# What Loss Functions Do Humans Optimize When They Perform Regression and Classification

**DOI:** 10.1101/2023.09.19.558376

**Authors:** Hansol X. Ryu, Manoj Srinivasan

## Abstract

Studying how humans perceive patterns in visually presented data is useful for understanding data-based decision-making and potentially understanding visually mediated sensorimotor control. We conducted experiments to examine how human subjects perform the simplest machine learning or statistical estimation tasks: linear regression and binary classification on 2D scatter plots. We used inverse optimization to infer the loss function humans optimize when they perform these tasks. Minimizing the sum of regression error raised to the power of 1.7 best-described human performing regression on sparse data. Loss functions with lower exponents, which are less sensitive to outliers, were better descriptors for regression tasks performed on less sparse data consisting of more data points. For the classification task, minimizing a logistic loss function was on average a better descriptor of human choices than an exponential loss function applied to only misclassified data. People changed their strategies as data density increased. These results represent overall trends across subjects and trials but there was large inter- and intra-subject variability in human choices. Future work may examine other loss function families and other tasks. Such understanding of human loss functions may inform the design of applications that interact with humans better and imitate humans more effectively.

## 1 Introduction

Studying how humans perceive patterns in visually presented noisy data is useful for understanding data-based decision-making (Keim et al., 2006; Moore, 2017; Kahneman et al., 2021), sensorimotor control under uncertainty due to disturbance and noise (Körding & Wolpert, 2004a; Srinivasan, 2009; Körding & Wolpert, 2004b; Todorov & Jordan, 2002), and more directly, visual perception (Glass & Pérez, 1973; Glass & Switkes, 1976; Dittrich, 1993). Examples of humans interpreting visually presented data include examining medical images (Lewandowsky & Spence, 1989), driving a vehicle (Hills, 1980; Baron & Kleinman, 1969), playing a sport (Davids et al., 2005), or living life in general (Palmer, 1975). On the simpler end of this spectrum of task and data complexity is interpreting 2D scatter plots of scientific data. Here, we study humans performing two simple types of visual pattern recognition tasks involving scatter plots, coinciding with fundamental machine learning and statistical inference problems: 1) linear regression (or “fitting”), that is, finding a curve that best represents continuous-valued input-output data (figure 1A); and 2) binary classification, that is, finding a decision boundary that separates pre-labeled data into two categories based on their labels (figure 1B).

**Figure 1:**
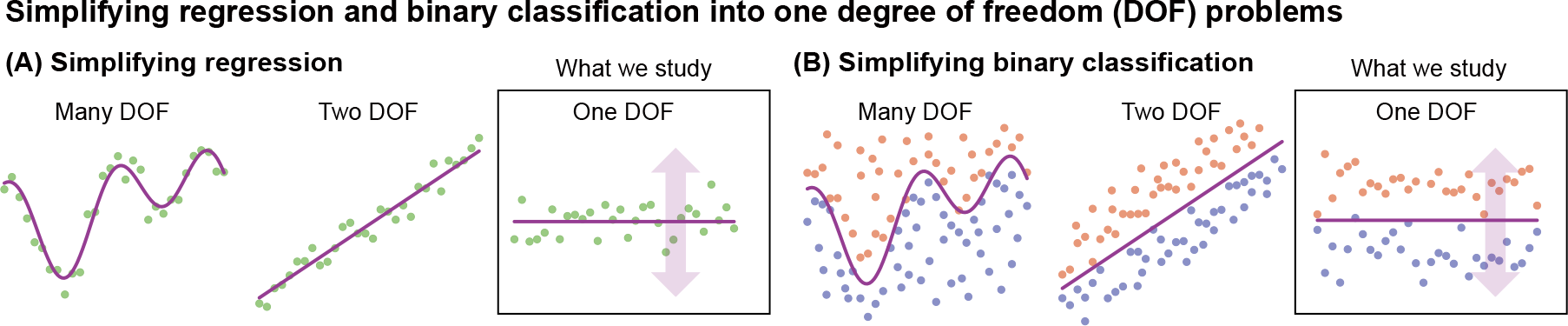
Simplifying regression and classification. A) Regression problems of decreasing complexity many degrees of freedom, two degrees of freedom, and one degree of freedom. We consider the simplest regression problem where the output function is a constant, so that there is only one degree of freedom as shown in the third panel. B) Classification problems of decreasing complexity: many degrees of freedom, two degrees of freedom, and one degree of freedom. Similarly, we consider the simplest binary classification problem where the classifier is a constant function, so there is only one degree of freedom. The one degree of freedom problem is sufficient to infer how prediction errors are penalized by the loss function, while deliberately ignoring issues such as model complexity, generalization, and over-fitting.

**Figure 2:**
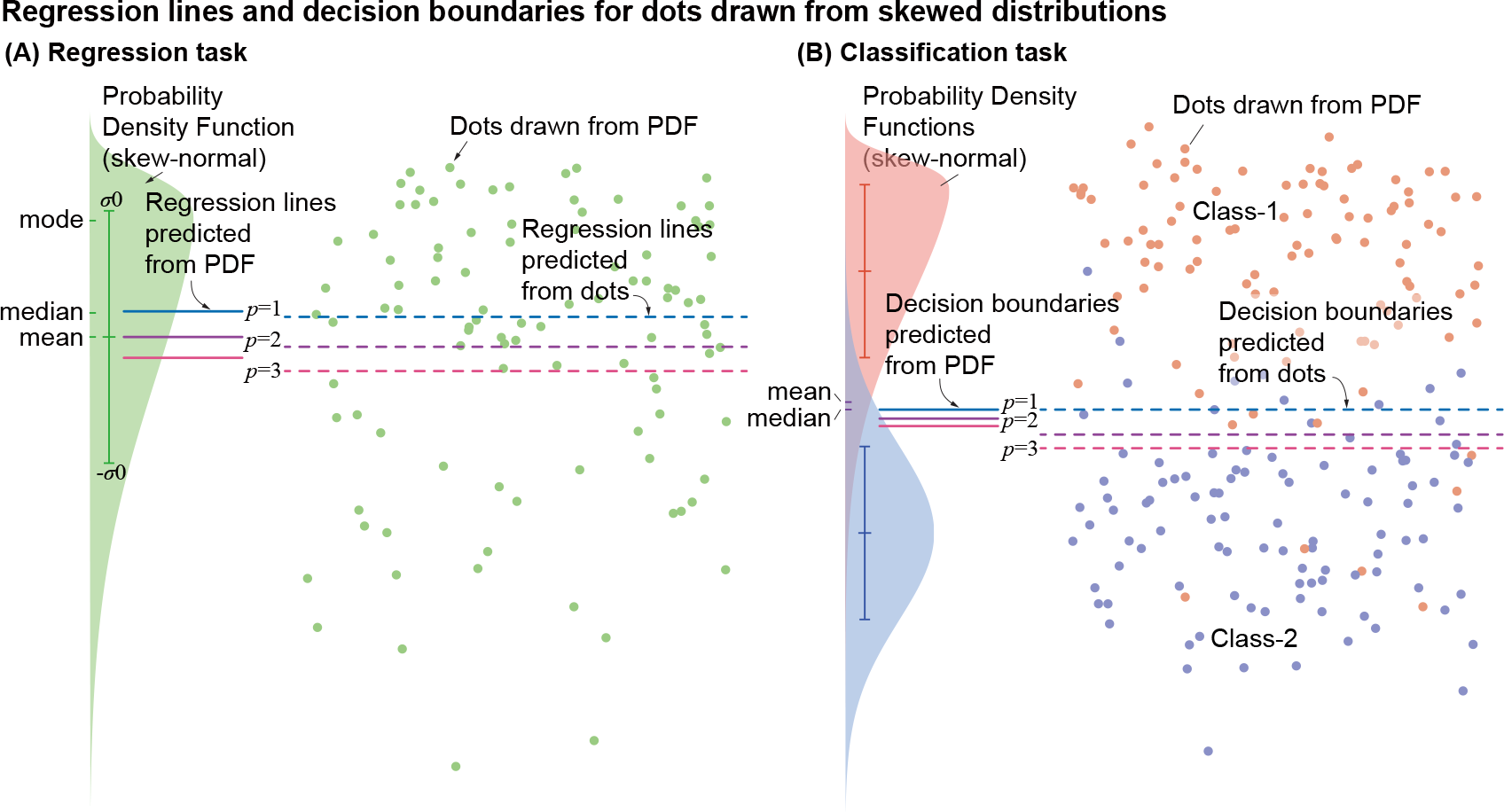
Regression lines and decision boundaries that minimized various objective functions on skewed probability distributions. (A) A skew-normal distribution (green shaded) was used to generate regression test dataset. (B) One skew-normal (orange shaded) and one normal distribution (blue shaded) were used to generate classification task test data. Probability distribution functions (PDFs, shown as shaded graphs) of skew-normal distributions have distinct mode, median, and mean. Error bars represent mean *±* variance of each PDF. Vertical locations of the dots were drawn from the PDFs. The location of the regression lines or decision boundaries depends on the exponent parameter *p* of the loss function they minimize. These locations calculated from the PDFs and from the actual dot locations are close to each other, but do not perfectly match when there are only a finite number of dots.

Classical machine learning algorithms for regression and classification usually find a solution that minimizes a loss function (Bishop & Nasrabadi, 2006). For regression problems, one commonly used loss function is the mean squared error (Luenberger, 1997; Bishop & Nasrabadi, 2006), equivalently, the *L*_2_ norm of the error, resulting in the ordinary least squares (OLS) regression. Other commonly used loss functions are mean absolute errors (equivalently, *L*_1_ norm) and the mean perpendicular distance between the regression surface and data (orthogonal regression). For binary classification problems, commonly used loss functions include cross-entropy, also called logistic loss function or log loss (Shore & Gray, 1982), and hinge loss function, used for support vector machines (Drucker et al., 1996). Loss functions are chosen for their favorable properties including robustness to outliers, over-fitting prevention, convexity, and uniqueness of the solution (Wang et al., 2022). Here, we examine whether human behavior in regression and binary classification tasks can be described as a solution of an optimization problem minimizing a loss function, and aim to characterize the loss function through experiments (figure 3). Different loss functions make different predictions on how humans would perform the tasks on each data (figure 2), and we use this fact to determine the loss function that best predicts observed human behavior, a process sometimes called inverse optimization (Tarantola, 2005; Liu et al., 2005).

**Figure 3:**
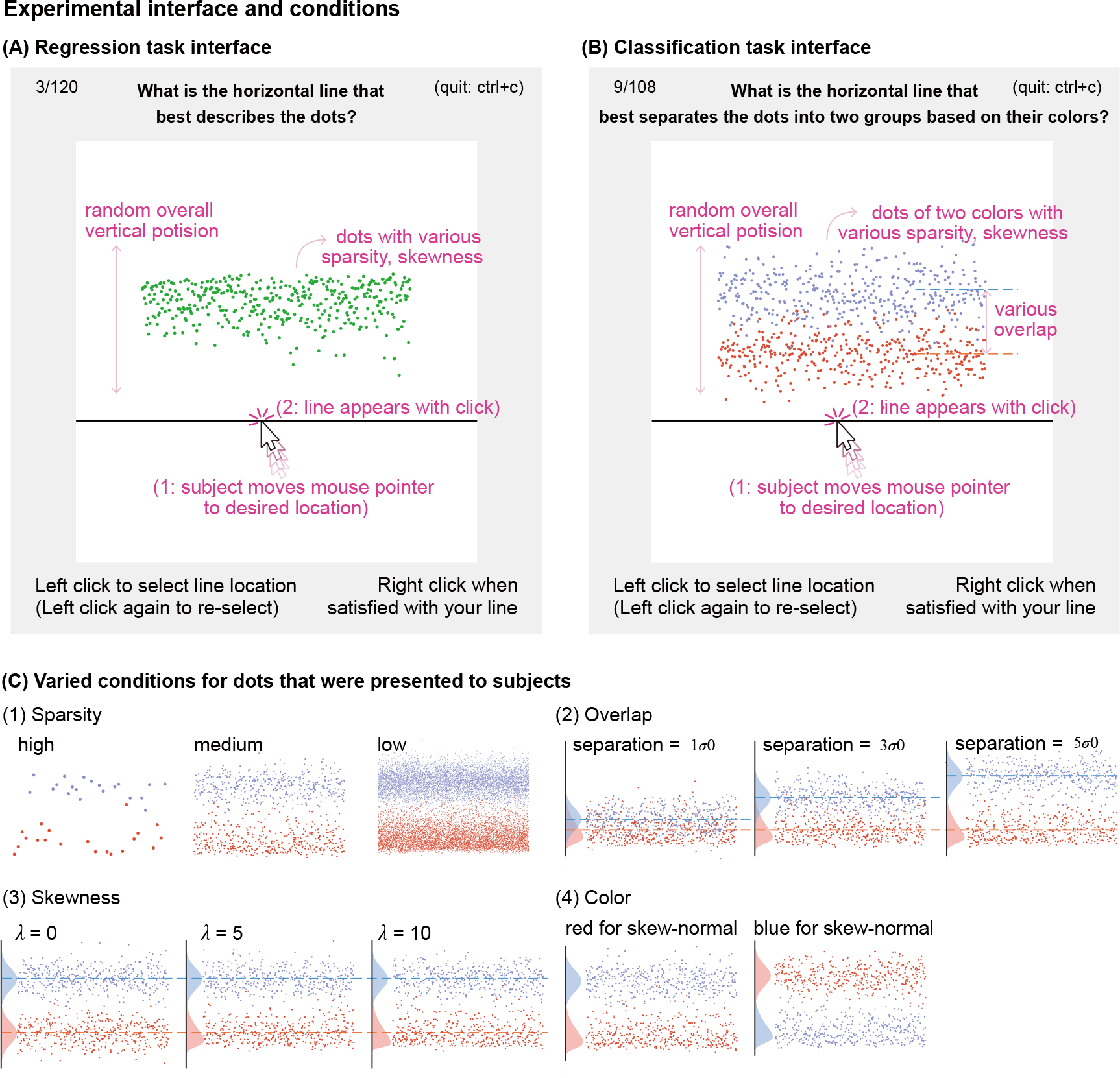
Testing interface for (A) regression and (B) classification tasks, and (C) conditions that were varied when dots were generated, to be used as testing datasets. On the testing interface (A and B), the task goal was shown on the top of the testing interface with the trial number and termination instruction on the side. Instructions on how to select a line and move to the next trial were shown on the bottom as text. Dots are shown in the white window of the test interface, and when a subject selects a line location by clicking the left button of the mouse, a horizontal line appears at that location. Subjects were free to select the line again as many times as they wanted. Pink labels are added only for the paper and are not shown to subjects, and some components have been re-sized here for illustrative purposes. The vertical locations of the dots were generated from probability distributions of varied conditions as shown in panel (C). Four conditions, (1) sparsity (2) overlap (3) skewness, and (4) color, were changed to generate testing data for classification tasks. The conditions used for the regression were subsets of those shown.

Some studies have attempted to characterize the strategies human subjects use in visual tasks (Lewandowsky & Spence, 1989), but most studies did not seek to provide quantitative descriptions of the loss function. For instance, it has been reported that human subjects were able to reject outliers when performing regression (Correll & Heer, 2017), sometimes better than some mathematical methods (Wainer & Thissen, 1979). For fitting a straight line to data, subjects selected the slope closer to that of the first principal component, instead of the least mean squares of vertical distances (Mosteller et al., 1981). Subjects with formal regression training selected lines closer to the least square fit, whereas subjects without regression training seemed to use other heuristics (Gillan, 2020). These studies compared human choices to the solutions of a few discrete strategies, rather than considering strategies on a continuum using inverse optimization approaches. These studies are also limited to particular visual datasets used and may have limited generalizability: for instance, the scale of the data axis was observed to affect human perception (Cleveland et al., 1982). Another study performed inverse optimization in a sensorimotor task for a narrow set of conditions (Körding & Wolpert, 2004a).

Relatively little has been done on inferring loss functions from human classification, as human labels are usually regarded as ground truth for machine learning and not analyzed independently. In one study, the support vector machine best predicted human behaviors among four algorithms considered for gender classification tasks on human facial images (Graf et al., 2006). Another study observed how subjects perceive transforming geometric shapes, describing when humans start to classify a transforming shape into a different shape (Gopsill et al., 2021). A study investigating the classification of scatter plot data reported high failure rates of about 50%, when evaluating an algorithm’s performance via manual inspections (Sedlmair et al., 2012). Thus, this study showed discrepancies between algorithmic output and human choices, but did not aim to characterize human behavior during classification tasks using inverse optimization.

In real-world applications, regression and classification problems can be high-dimensional in terms of both data dimensionality and model complexity (such as via neural networks). Related issues such as model complexity, generalization, and over-fitting prevention need to be considered, in order to decide which optimization algorithm will be used to solve a problem. Here, we intentionally ignore these latter issues by considering the simplest one degree of freedom regression and binary classification problems (figure 1), which are minimally sufficient to infer how humans penalize errors of different magnitudes via minimizing a loss function. Secondly, we examine whether human perception of the pattern depends on data sparsity. Many computer algorithms are not designed to change the loss function based on the amount of data, but humans may perceive different data amounts as qualitatively different. Classifying a sparse, handful of dots into two groups could appear to be different from classifying thousands of dots into two groups, which resembles identifying and separating two visually distinct areas in a digital image. Here, we inferred regression and classification loss functions from human experiments with different visual data sparsity, showing how visual sparsity can affect behaviors.

## 2 Methods

### 2.1 Experiments

Subjects participated with informed consent, and the experimental protocol was approved by the Ohio State University Institutional Review Board. Subjects performed two types of tasks: 1) a regression task (*N*_subjects_ = 23 subjects) and 2) a binary classification task (*N*_subjects_ = 23 subjects; one subject’s medium sparsity data and one subject’s low sparsity data was corrupted, thus excluded). For regression, we showed dots of a single color on a computer screen, and asked “What is the horizontal line that best describes the dots?” (figure 3A). For classification, we showed dots of two colors, with various amounts of overlapping, and asked “What is the horizontal line that best separates the dots into two groups based on their colors?” (figure 3B). Subjects used either a keyboard or mouse to select the vertical location of the horizontal line that best fit or classified the dots. Subjects could re-select the lines as many times as they wanted without a time limit. Subjects performed the test either on the experimenter’s computer or on their own. After the test, subjects optionally filled out a questionnaire about what their strategies were for each sparsity level and whether they were familiar with regression or classification algorithms.

Subjects performed tests with data of different sparsity and skewness, drawn from specific probability distributions (figure 3C). There were 3 sparsity conditions, and 40 and 36 trials per sparsity condition for regression and classification tasks, resulting in 120 and 108 trials total per subject. About two-thirds of the subjects (*N*_subjects_ = 16 for both tasks) were tested with a partially randomized order: they performed the task on high sparsity data first, then medium, and then the low sparsity, while skewness and other conditions within each sparsity condition were randomized. Other subjects were tested in a fully randomized order including the sparsity. We made this decision to keep most of the subjects unaware of the underlying probability density distribution when they perform a task on high-sparsity data.

### 2.2 Probability distributions for the testing data

#### 2.2.1 Datasets for regression task

We drew testing data from skewed probability distributions, so that minimizing different loss functions predicts different regression lines (figure 2). We used skew-normal distributions *𝒮 𝒩* (*λ*) given by:

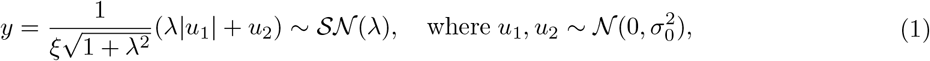

when *λ* is a shape parameter (Azzalini, 1985; Henze, 1986). Shape parameter *λ* = 0 gives normal distribution 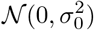 with a mean 0 and variance 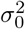, and increasing |*λ*| increases skewness. The normalization constant 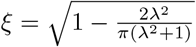ensures that the variance remains 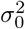 regardless of *λ*. We used shape parameter values *λ* = 0, *±* 5, *±* 10, *±* 40 (figure 3C-3). For the skew-symmetric distributions (*λ* ≠ 0), the mode is at the head of the distribution where probability density is the highest, and the median and the mean were located on the longer tail (figure 2). When the distribution is symmetric and unimodal (*λ* = 0), regression using typical loss functions mostly yields the same solution, as the mean, median and mode are all identical. We still included *λ* = 0 trials, but trials with *λ* ≠ 0 provide more information on human loss functions.

#### 2.2.2 Datasets for classification task

We generated dots of two colors for the binary classification task, one from a normal distribution and the other from a skew-normal distribution, so that different loss functions yield different classification boundaries (figure 2). We use the same skew-normal distributions as for the regression task with *λ* = 0, *±*5, *±*10 (figure 3C-3). We changed the overlap between two distributions, separating the theoretical means of the two distributions by *σ*_0_, 3 *σ*_0_, 5 *σ*_0_ (figure 3C-2). The normal distribution (*λ* = 0) was always on the skew-normal distribution’s longer-tail side and ensured overlap of the two sets: we re-generated the data until the lowest dot of the upper distribution was lower than the highest of the lower distribution by 0.05 *σ*_0_. We included *λ* = 0 trials, but *λ* ≠ 0 trials, when different loss functions have different predictions, are more informative.

#### 2.2.3 Sparsity

For both regression and classification tasks, we tested high-, medium-, and low-sparsity datasets, respectively, using *N*_data_ = 20, 400, 8000 dots of each color (figure 3C-1). Only the vertical positions of the dots were randomly drawn from the specified distributions; the horizontal positions were evenly spaced. The size of the dots was scaled across the different sparsity conditions such that the area of each dot multiplied by the number of dots stayed the same. We changed the overall vertical position of the dots on the display each trial to minimize the effect of the previous trial. We used orange and blue colors for the classification task, which are usually distinguishable even with some color vision deficiency (Wong, 2011). We randomized the vertical order of the normal and skew-normal distribution, and the color of the top versus bottom distribution (figure 3C-4).

### 2.3 Loss function models to predict regression lines and classification decision boundaries

#### 2.3.1 Loss functions for regression

The loss function *J*_*μ*_(*ŷ*, ***y***) is a function of the data points ***y***, a candidate vertical location of the regression line *ŷ*, and possibly other hyperparameters *μ*, such that the optimal regression line location *ŷ*^*\**^ is obtained by minimizing this loss function as follows:

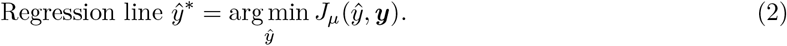

We may denote *ŷ*^*\**^ as *ŷŷ*^*\**^(*J*_*μ*_, ***y***) to show dependence on both the loss function *J*_*μ*_ and the data ***y***. In this study, we considered loss functions that are sums of the absolute regression error raised to various exponents *p* (figure 4A-1). Regression error here refers to the difference between the regression line and each data point. When the vertical location of the *i*-th dot is ***y***(*i*), and the regression error of the dot is ***x***(*i*) = ***y***(*i*) − *ŷ*, the regression loss function with exponent parameter *p* is given by:

**Figure 4:**
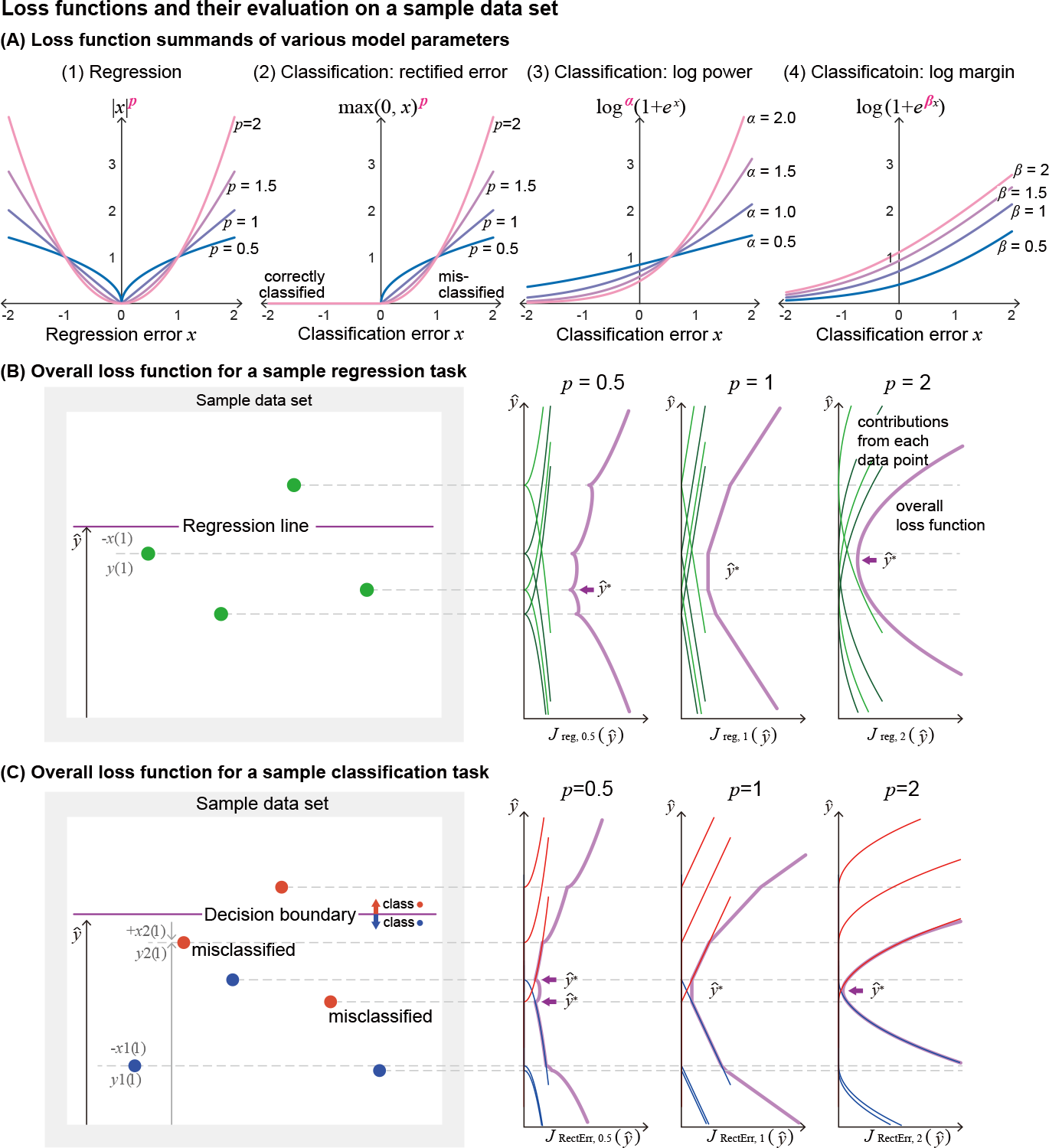
Demonstration of various loss functions for regression and classification tasks. (A) Loss function summands of various parameters, as functions of (1) regression error *x* for regression task, or functions of (2-4) classification error *x* for classification task. (1) Regression loss function with exponent parameter *p*. (2) Rectified error loss function, with exponent parameter *p*. (3) Log power loss function with power parameter *α*. (4) Log margin loss function with margin parameter *β*. (B) Evaluation of overall regression loss for a sample regression task dataset ***y***, as a function of the regression line location *ŷ*, with various exponent parameters. The overall loss (purple thick line) is a sum of contributions from each data point (green thin lines). (C) Evaluation of overall rectified error loss for a sample classification task dataset ***y***_1_, ***y***_2_, as a function of the regression line location *ŷ*, with various exponent parameters. The overall loss (purple thick line) is a sum of contributions from orange class (orange thin lines) and blue class (blue thin lines) data points. Orange dots have positive classification errors when the decision boundary is above them (two orange dots marked as “misclassified” have positive losses for the decision boundary shown in the illustration), and blue dots have positive classification errors when the decision boundary is below them. *ŷ*^*\**^ indicates global minima.

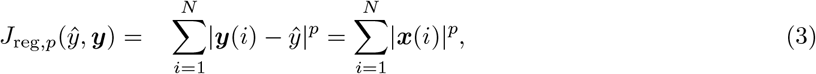

equivalent to using the error vector’s *L*_*p*_ norm. The individual summand in the loss function (figure 4A-1), |*x*| ^*p*^, defines how error from each point contributes to the total loss (figure 4B). It is symmetric with respect to zero error, so that error on either side of the regression line has the same loss.

The effect of the exponent *p* on the overall loss could be understood in terms of how larger regression errors are penalized relative to the small errors. For example, regression loss function summand |*x*| ^2^ grows faster than |*x*| ^1^, thus penalizing larger error relatively more (figure 4A-1). For example, minimizing *J*_reg,2_ results in a regression line more towards the tail of the distribution compared to minimizing *J*_reg,1_ on a skew-normal distribution (figure 2A), because moving the regression line towards the tail of the distribution reduces larger errors that come from the further points.

#### 2.3.2 Loss functions for binary classification

The loss function for binary classification *J*_*μ*_(*ŷ*, ***y***_1_, ***y***_2_) is a function of the candidate decision boundary location *ŷ* and the data points ***y***_1_ and ***y***_2_, belonging to the two classes (two colors), class-1 and class-2 respectively. The decision boundary divides the data points into two groups, so that data below the boundary is classified as class-1 and data above the boundary is classified as class-2. The optimal decision boundary *ŷ* ^*\**^ is defined as minimizing the loss function *J*_*μ*_ (figure 4C) as follows:

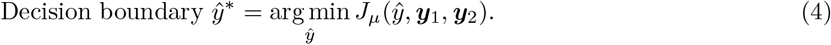

We may denote *ŷ*^*\**^ as *ŷ*^*\**^(*J*_*μ*_, ***y***_1_, ***y***_2_) to show dependence on both the loss function and the two datasets.

Loss functions for classification are defined such that misclassified points increase the loss more than correctly classified points do (Bishop & Nasrabadi, 2006). We consider two types of loss functions: those that only penalize misclassified points and are not affected by correctly classified points, and those that also account for correctly classified points. Our datasets were not linearly separable and always had misclassified points. Let the location of the *i*-th member of the two classes be ***y***_1_(*i*) and ***y***_2_(*i*). Members of ***y***_1_ but located above a candidate decision boundary *ŷ* (i.e., ***y***_1_(*i*) − *ŷ>* 0), and members of ***y***_2_ but located below the boundary (i.e., *ŷ* ***y***_2_(*i*) *>* 0), are misclassified by the decision boundary *yŷ*. We define classification error ***x***_*c*_ for for *c* = 1, 2 as follows: for data points in class-1, ***x***_1_(*i*) = ***y***_1_(*i*) − *ŷ* and for data points in class-2, ***x***_2_(*i*) = *ŷ* − ***y***_2_(*i*), ensuring that the classification error ***x***_*c*_(*i*) is positive when misclassified. We now describe specific loss functions that we considered.

#### Rectified error classification loss function

As an example of a loss function that only penalizes misclassified points, we investigated the effect of the exponents on the positive part of the classification error (“rectified error”, figure 4A-2). We define the rectified error classification loss function with exponent parameter *p* as:

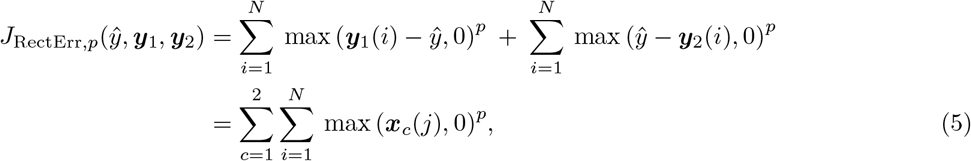

where the positive part or rectification function max(*x*, 0) is the ReLU function. The exponent parameter *p* has a similar effect to that in the regression loss function. A loss function with bigger *p* penalizes bigger errors relatively more than a loss function with smaller *p* does. For distributions used here, since the skew-normal distribution has a heavier tail than a normal distribution on the overlapping side, decision boundaries tend to move towards the tail of the skew-normal distribution when the exponent *p* of the loss function increases (figure 2B). When *p* = 1, the decision boundary divides data points such that there are equal numbers of misclassified points on both sides, analogous to a median.

#### Log power classification loss function and log margin classification loss function

When correctly classified points also contribute to the loss function, the expected effect is that the distance between the correctly classified points and the decision boundary increases. This gives a better margin for the decision boundary, thus the classification becomes “more correct”. However, the loss still needs to increase more with misclassified points, in order to have sensible classification. In this study, we used two one-parameter families of functions derived from the logistic loss function, in which loss smoothly increases with positive classification error (from misclassified data) and decreases with negative error (from correctly classified data). The “Log power” classification loss function with power parameter *α* is given by:

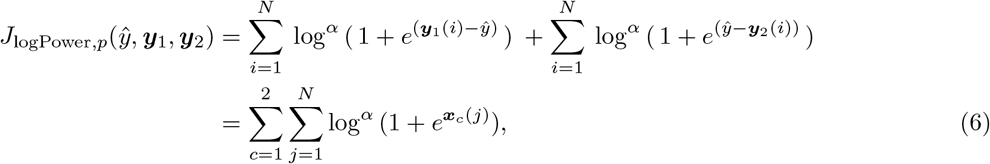

and the “log margin” classification loss function with margin parameter *β* is given by:

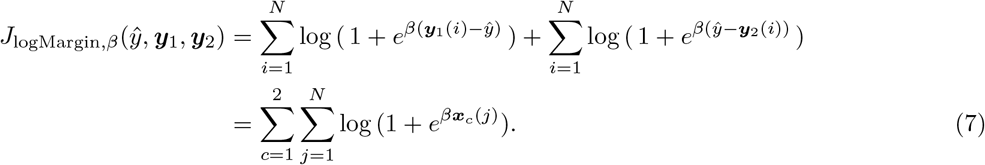

We analyzed the effect of the exponent in the”log power” loss (figure 4A-3) and the margin parameter in the “log margin” loss (figure 4A-4, Masnadi-Shirazi & Vasconcelos (2015)). In addition to these one-parameter families of cost functions, we considered two standard cost functions: the hinge loss for support vector machines and the logistic or cross-entropy loss function of logistic regression (Wang et al., 2022).

### 2.3.3 Inverse optimization to infer loss functions from subjects’ responses

To evaluate which loss function best describes each subject’s responses, we minimized the mean squared distance between the subject’s responses and model-predicted regression lines or decision boundaries for each trial, averaged across all trials. Let the model-predicted regression line (eq. equation 2) or decision boundary (eq. equation 2) on the *k*-th trial be 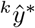. Let the subject’s response on the *k*-th trial be ^*k*^*γ*. Then, the root mean squared error (RMSE) between human response and model prediction is:

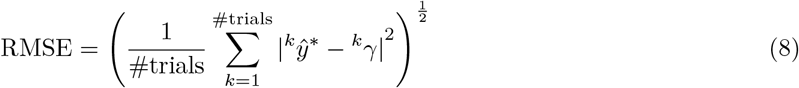

For each trial, the model prediction 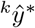 depends on the loss function *J*_*μ*_ used, the hyperparameters *μ* of the loss function, and the particular dataset from the *k*-th trial. So, we may denote the mean squared error in eq. equation 8 as RMSE(*μ*), focusing on the dependence on the loss function hyperparameters *μ*, suppressing the dependence on the datasets used and the particular loss function *J*_*μ*_ for simplicity. For the four parameterized loss functions considered, the hyperparameters *μ* are *p* for the regression loss function, and *p, α*, and *β*, respectively, for the three classification loss functions. Within a given one-parameter family of loss functions *J*_*μ*_, the best-describing loss function is determined by computing the best-describing loss function parameter *μ*^*\**^ as minimizing RMSE(*μ*):

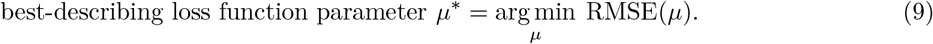

We used a quasi-newton algorithm (MATLAB fmincon) to find the location *ŷ*^*\**^ for most of the cases, as the loss function usually has a unique global minimum without other local minima. The exceptions were for exponents *p* equal to or smaller than 1 (*p <*= 1 in eq. equation 3 and eq. equation 5), as there could be multiple local minima. We used *p* = 1.001 as a proxy for *p* = 1, as *p* = 1 could produce infinitely many minima. For *p <* 1, we evaluated the loss function at the testing data point locations, as the minima will appear at those places (figure 4B and C, *p <* 1). When we found two global minima with identical values (*p <* 1 in classification), we picked the model prediction *ŷ*^*\**^ to be the minimum closer to the subject selection.

## 3 Results

### 3.1 Regression: subjects effectively track measures of central tendency

Subjects effectively tracked standard measures of central tendency of the data shown in each trial (figure 5). Specifically, the location of the mean, the median, and other measures of central tendency changed substantially between trials, and the locations of the mean and the median were highly predictive of where the subjects clicked. The linear model from the mean of the distribution to where the subjects clicked had 97.1% R^2^ value and the median of the distribution to where the subjects clicked had 97.3% R^2^ value, when averaged across all subjects and all trials. Thus, subjects are able to track the overall location of the distribution well. The rest of the results section is about the small but significant differences between how well different measures of central tendency, or equivalently, minimizing different loss functions, predict the subject responses.

**Figure 5:**
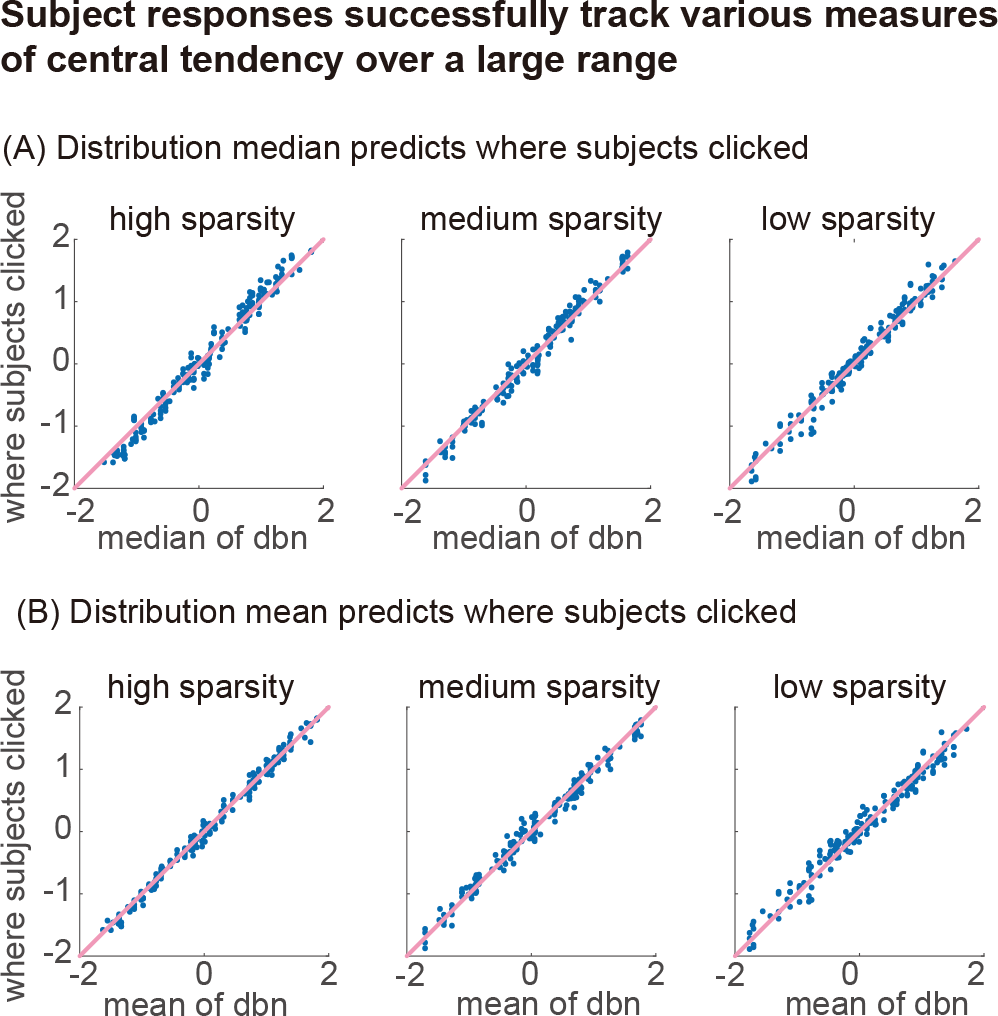
Subject responses compared to various centers of the distributions. Subjects effectively tracked standard measures of central tendency. The location of the median and the mean varied substantially on the screen, and subjects systematically tracked these.

### 3.2 Regression: loss function exponent reduces with reducing sparsity

The RMSE (eq. equation 8) between the subjects’ responses and the model-predicted regression lines, averaged across all subjects and all trials for a given sparsity, has a U-shaped curve with respect to the *L*_*p*_ exponent *p* (figure 6 left). We obtained the best describing exponent *p*^*\**^ by minimizing the overall RMSE for each sparsity. The best-describing parameters were different for different sparsity levels. We found that *x*^1.7^ for high sparsity (20 dots), *x*^1.2^ for medium sparsity (400 dots), *x*^0.7^ for low sparsity (8000 dots) best predicted human regression lines on average. Thus, as sparsity decreased, or in other words, as the data density increased, subjects tended to choose regression lines described by a loss function with a smaller exponent.

**Figure 6:**
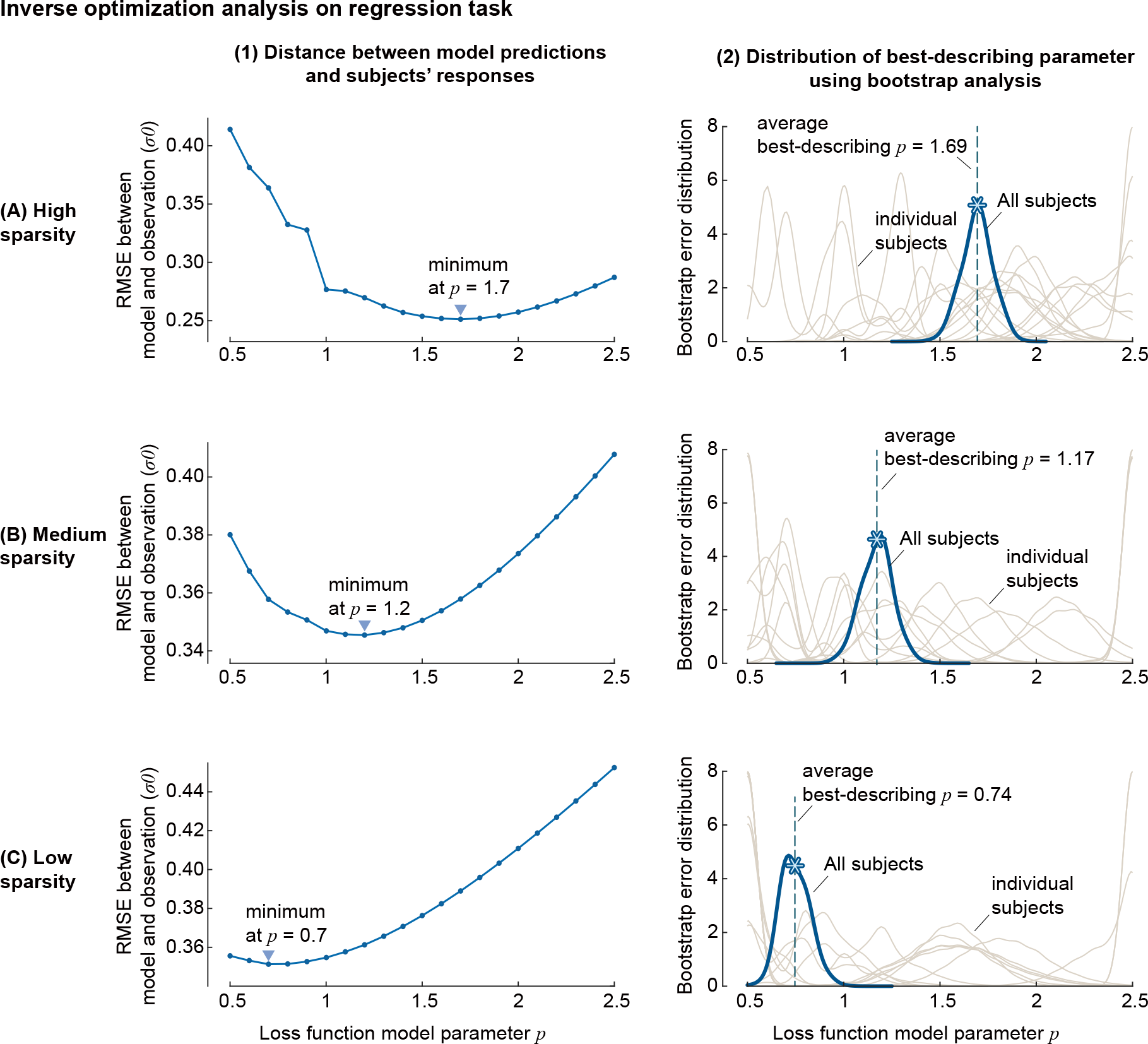
Comparison of subjects’ responses to the predicted regression lines. Regression lines were predicted using loss function models of various exponent parameter *p*, for different sparsity levels (top to bottom). On the left are the RMSE between the predicted regression lines and the subject’s response, normalized by PDF variance *σ*_0_, and the downward triangle indicated the loss function parameters that had minimum RMSE for the given sparsity. On the right are the distributions of best-describing loss function parameters inferred from the RMSE, using bootstrap method. Thin lines indicate best-describing loss function parameter for each subject, and the thick black line shows that of the entire subjects. The asterisk on the tick line indicates the average of the best-describing loss function parameter from each bootstrap sample.

To compute the uncertainty in these estimates, we performed bootstrap resampling of trials from all subjects and all trials of a given sparsity and recomputed the best-describing exponent *p*^*\**^ for each sample to obtain bootstrap-based error distributions for *p*^*\**^ (figure 6 right). The average best-describing loss function parameters obtained from this bootstrap analysis were 1.69, 1.17, 0.74 for high, medium, low sparsity, with relatively clear peaks at those parameter values, and error standard deviations of about 0.15 for each sparsity. Right-tailed paired t-test had *p* values of 0.0032 between high- and medium-sparsity, and 0.0004 between medium- and low-sparsity, suggesting that best-describing parameters were significantly lower as sparsity decreased. We note, however, that the RMSE landscape near *p*^*\**^ is quite flat, that is, has low curvature (figure 6 left), indicating that substantially different exponents predict a small increase in error, partly an indication of inter- and intra-subject variability in responses, as described later.

### 3.3 Classification: rectified error loss had worst performance and exponent changed with sparsity

RMSE using rectified error loss function had the worst performance among the loss functions we considered (figure 7 left). Commonly used loss functions, logistic regression and support vector machines (SVM) were closer to subjects’ responses than rectified error, with SVM being closer than logistic regression. The other two parameterized loss functions were also systematically better. This provides limited evidence that subjects do not just consider error in misclassified points while performing the classification task, as rectified error loss function is the only one of our loss functions that only used misclassified points for computing loss.

**Figure 7:**
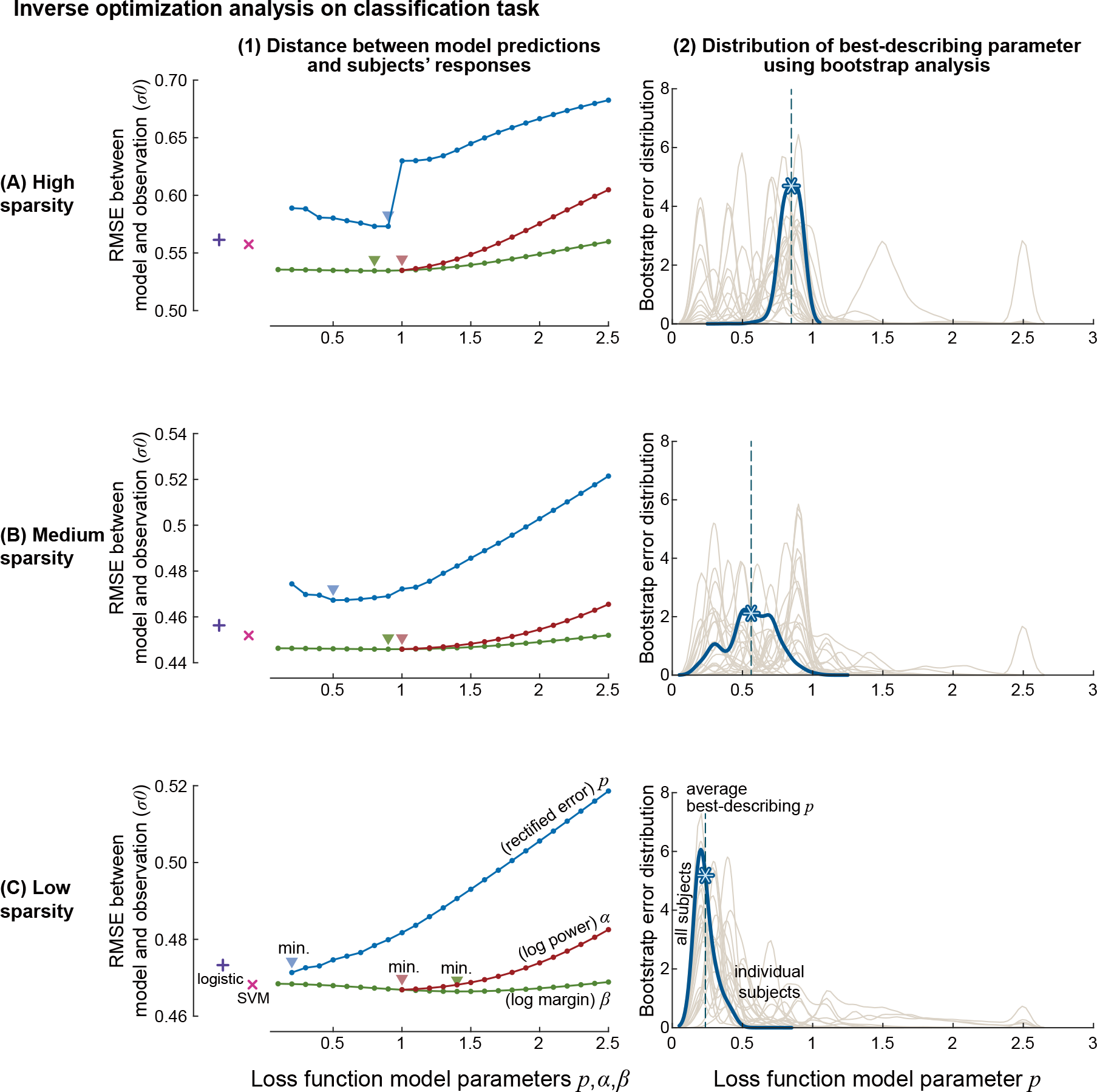
Comparison of subjects’ responses to the predicted decision boundaries. Decision boundaries were predicted using rectified error (blue), log power (red), and log margin loss functions (red), with their model parameter *p, α, β*, for different sparsity levels (top to bottom). Errors from the decision boundaries obtained from logistic regression classifier (plus mark) and SVM (x mark) are also shown as a reference. On the left are the RMSE between the subject’s response and the decision boundaries obtained by minimizing loss functions, normalized by PDF variance *σ*_0_, and the downward triangle indicated the loss function parameters that had minimum RMSE for the given sparsity. On the right are the distributions of best-describing exponent parameter *p* for rectified error loss function, inferred from the RMSE, using bootstrap method. Thin lines indicate best-describing loss function parameter for each subject, and the thick black line shows that of the entire subjects. The asterisk on the tick line indicated the average of the best-describing loss function parameters from each bootstrap samples.

RMSE using rectified error loss function averaged across all subjects and all trials had minima within the parameter range we searched, and the best-describing parameter changed systematically with the sparsity of the dots (figure 7 left). Rectified error to the power of *p*^*\**^ = 0.9, 0.5, 0.2 were the best descriptors of subjects’ responses for each sparsity using rectified error loss functions. Our observation of smaller best-describing *p*^*\**^ for lower sparsity roughly means that subjects chose decision boundaries closer towards the skew-symmetric distribution when sparsity decreased; in other words, more dots from the skew-symmetric distribution tended to be misclassified than dots from the symmetric distribution as sparsity decreased.

Best-describing parameters obtained from bootstrap analysis had similar results. Best-describing parameters for rectified error loss function were *p* = 0.85, 0.75, 0.23 (figure 7 right) with standard deviations of about 0.15, 0.33 and 0.15 respectively. Right-tailed paired t-test had *p* values of 0.23 between high- and medium-sparsity, and 0.00037 between medium- and low-sparsity, suggesting that best-describing parameters of low-sparsity data were significantly lower than other high- and medium-sparsity data. Performing subject-specific analysis, we find that most of the subjects had best-describing exponent *p*^*\**^ lower than 1, which in general means that they drew the decision boundary closer to the skew-symmetric distribution, and more so as the sparsity decreased.

The RMSE curve for rectified error loss function at high sparsity had a distinct jump between *p* = 0.9 and 1.001 (figure 7 left), because *p <*= 1 could have multiple global minima (as seen in figure 4C *p* = 0.5 and 1) and we used the one that is closer to the subjects’ response for the analysis. Such discrepancy also exists in medium and low sparsity, but the effect is less noticeable because multiple minima, which can only exist between adjacent dots, are closer to each other as sparsity decreases.

### 3.4 Classification: log margin loss function best describes human responses

We considered two one-parameter families of loss functions — sum of log power (eq. equation 6), sum of log margin (eq. equation 7) — that considered error in both correctly classified and misclassified points. Their hyperparameters were an exponent parameter *α* and margin parameter *β* respectively. Log margin and log power in general had smaller RMSEs and, thus were better descriptors of subjects’ behavior than rectified error loss function, while log margin was the best descriptor among the three functions in the range we considered. Log margin at its minima also yielded decision boundaries closer to subjects’ behaviors than SVM and logistic regression did.

Log margin loss function model had minima within the parameter range we searched, and the best-describing parameter changed systematically with the sparsity of the dots (figure 7 left). Specifically, increased margin logistic function with margin parameter *β*^*\**^ = 0.8, 0.9, 1.4 were the best descriptors for each sparsity (figure 7 left), although RMSE curve is shallow for the wide range of the parameter for this loss function. The RMSE using log power had minima at *α*^*\**^ = 1.001 for all three sparsity conditions. This *α*^*\**^ = 1.001 was at one extreme of our evaluation range, as we did not evaluate values below *α* = 1.001 (a proxy for *α* = 1.0).

Best-describing parameters obtained from bootstrap analysis had similar results. Parameter for log power loss function *α*^*\**^ = 1.00, 1.04, 1.05 (Figure 8 left) and parameter for log margin loss function *β*^*\**^ =0.74, 0.94, 1.46 (Figure 8 right) were best-describing parameters obtained from the bootstrap analysis, with standard deviations of about 0.35, 0.6, and 0.7 respectively. Left-tailed paired t-test had *p* values of 0.0095 and 0.0021 between high- and medium-sparsity, and 0.738 and 0.059 between medium- and low-sparsity for log power and log margin loss function, suggesting that best-describing parameters of high-sparsity data were significantly lower than medium- and low-sparsity data.

**Figure 8:**
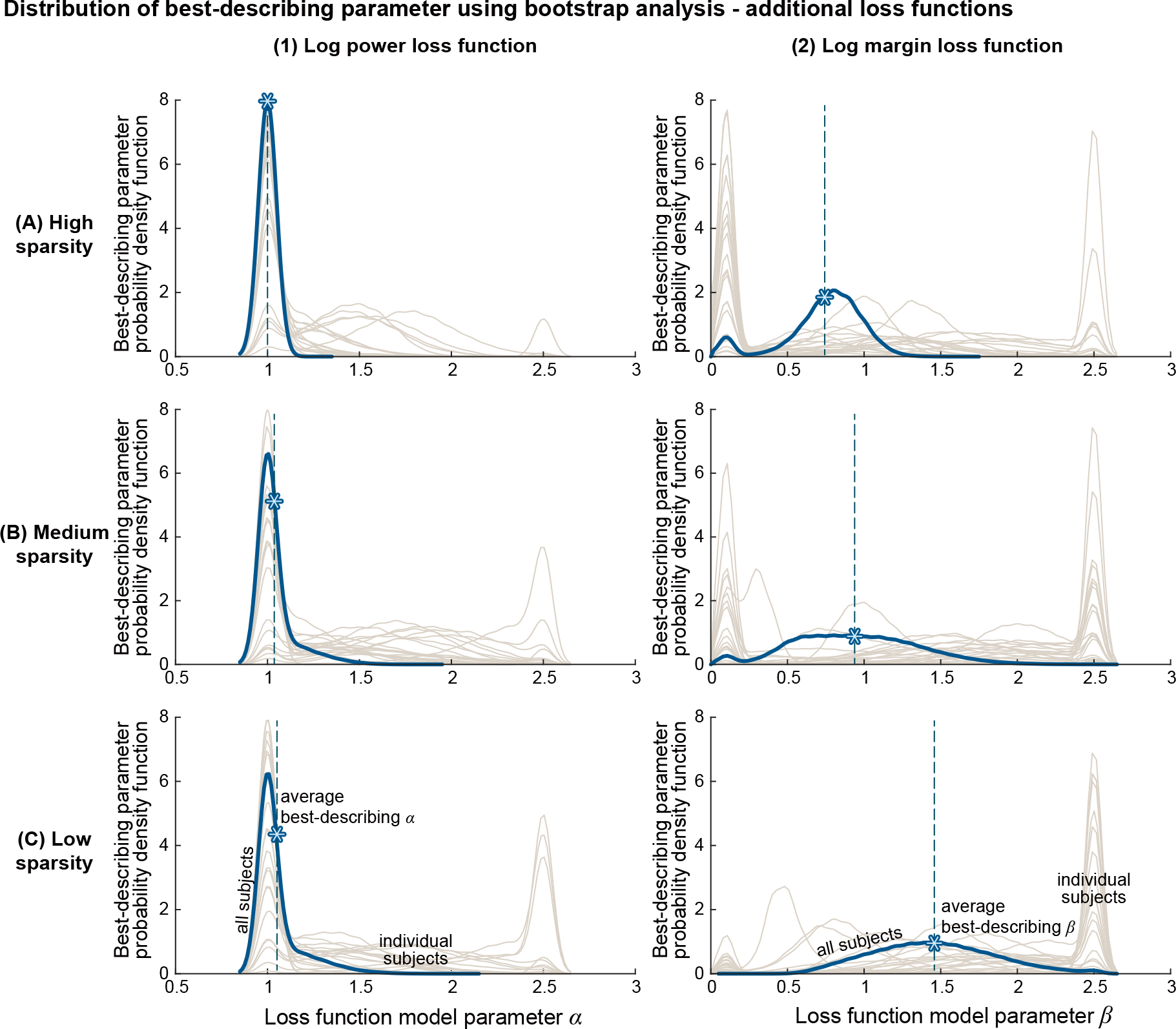
Best-describing parameters for log power and log margin loss function. Distributions of best-describing exponent parameter *α* and *β* were inferred from the RMSE by applying bootstrap method. Thin lines indicate best-describing loss function parameter for each subject, and the thick black line shows that of the entire subjects. The asterisk on the tick line indicates the average of the best-describing loss function parameters from each bootstrap sample.

### 3.5 Regression and classification: large inter-subject variabilities in responses and in loss functions

We obtained the best-describing regression and classification loss function hyperparameters for each subject and constructed a subject-specific bootstrap distribution of the best-describing hyperparameters. We found substantial inter-subject variability — seen in the wide ranges of peak locations among different thin lines in figures 6-7 (right), indicating best-describing parameters for individual subjects were not consistent across them. Furthermore, when we analyze their behavioral variability, the results also indicate substantial intra-subject variability — evident in the wide ranges covered by each of the thin lines in figures 6-7 (right), indicating best-describing parameters were not highly consistent across trials within the same subject. Parameters for rectified error in the classification task had fairly large inter- and intra-subject variability, although it was smaller than the variability observed in the regression task.

To more directly observe the inter-subject variability, a subset of our subjects (9 for the regression task, 13 for the classification task) were tested with the exact same arrangements of dots. Subjects’ responses (figure 9 thin lines) varied to a large degree, often more than the range of regression lines that are calculated by minimizing loss functions of various parameters (figure 9 thick lines). Even for the dots that were generated from symmetric probability density function(s) (figure 9 middle rows), where regression lines calculated using various loss functions lie close to each other at the middle of the distribution(s), (thicker lines on top of each other), subjects’ responses varied to a large degree.

**Figure 9:**
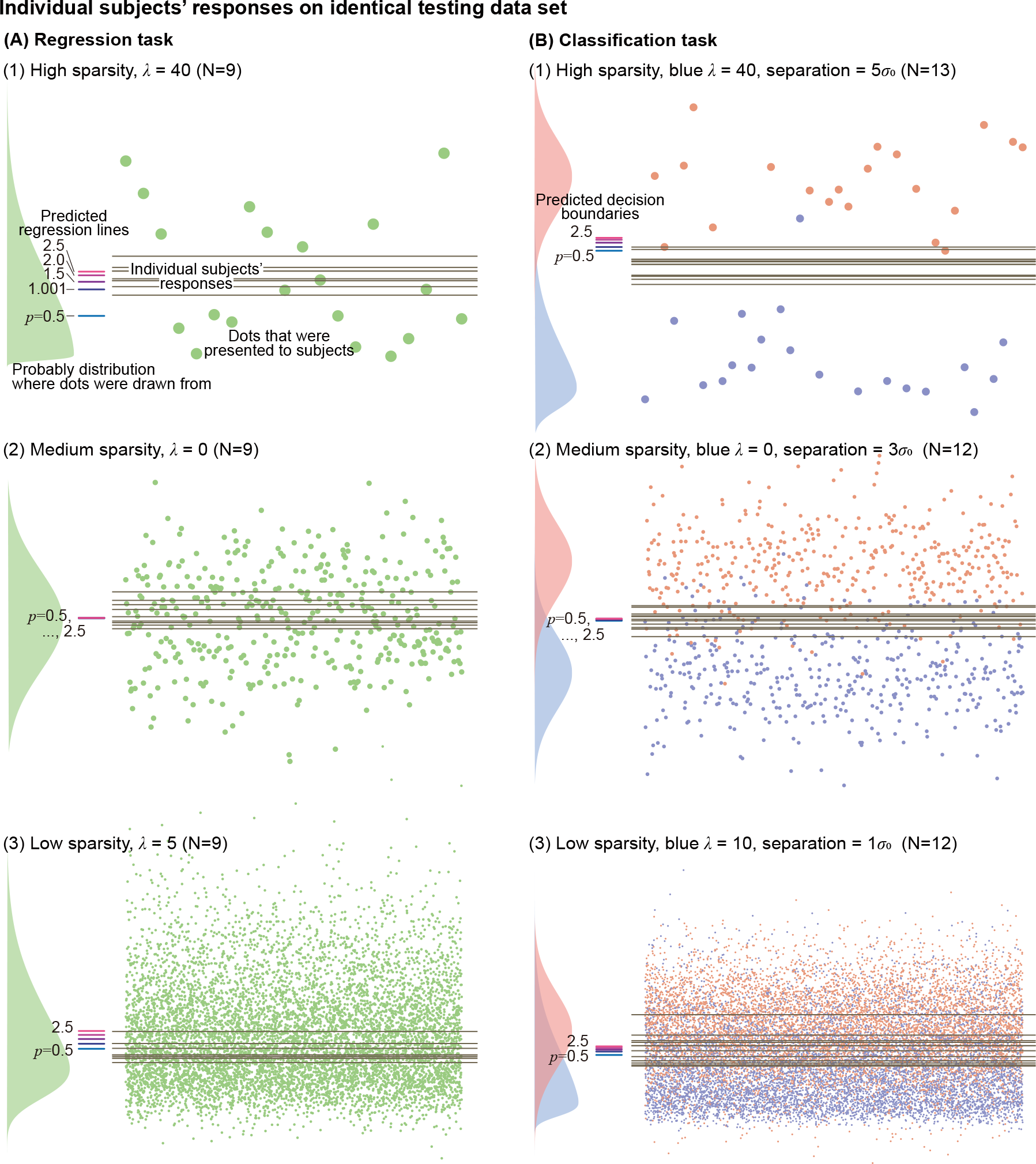
Various responses from multiple subjects on identical testing datasets. Responses from (A) regression task and (B) classification task, from representative trials of various conditions. The dots on each panel were drawn from the PDFs(s) shown on the left side of the panel. Regression lines that minimize regression loss functions of various exponent parameter *p*, and decision boundaries that minimize rectified loss functions of various exponent parameter *p* are indicated with the thick, short horizontal lines as references. The longer, thin lines on top of the dots are the responses from each subject.

### 3.6 Regression and classification: a weak trend of longer time for sparser trials

Subjects (*N*_subjects_ = 7) who were tested with fully randomized orders for regression task took 6.4*±* 20.3, 4.9*±* 10.9, 3.8*±* 4.2 (average*±* standard deviation) seconds to complete a regression task trial for high, medium, low sparsity. Both the mean and standard deviation of the completion time decreased on average with decreasing sparsity. Subjects (*N*_subjects_ = 7) who were tested with fully randomized orders for the classification task took 4.0 *±* 6.0, 3.7 *±* 8.8, 3.6*±* 9.1 seconds for high, medium, low sparsity. The average completion time slightly decreased with decreasing sparsity, but the standard deviation increased. We did not perform explicit statistical significance tests.

## 4 Discussion

We applied inverse optimization analysis on humans performing regression and classification tasks in visually presented data. For the regression task, minimizing the sum of error raised to the power of 1.69, 1.17, 0.74 were the best descriptions of the average subjects’ responses for high, medium, and low sparsity data. For the classification task, minimizing the sum of increased margin logistic classification error was a better descriptor among various types of loss functions we considered, with its best-describing margin parameters *β* being 0.74, 0.94, 1.46 for high, medium, low sparsity data. Among the loss function models that sum error powers of only misclassified data, exponents of *p* = 0.85, 0.75, 0.23 were the best descriptors, while logistic error functions that also take into account correctly classified data were in general better. These changes to the best-describing loss functions with decreasing sparsity had a common tendency: the regression line or decision boundary subject chose moved towards the mode of the skew-symmetric distribution, or in other words, towards the densest part of the dataset, as the data became denser.

These results qualitatively agree with some previous findings. A study that tried to infer a human loss function from a pointing task (Körding & Wolpert, 2004a) reported that minimizing the sum of distances raised to the power of 1.69 was the best descriptor of the average subjects’ behavior, when subjects observed about 60 data points during each trial. Our trials with the sparsest data consisted of 20 data points, and the best-describing loss function for the regression task was distance raised to 1.69, which is a close match with the previous finding. This exponent of 1.69 is also reasonably close to 2, which is used for the least mean squared method, so is in alignment with the study (Gillan, 2020) reporting that students with statistics training were more likely to choose a regression line based on least square method among other specific alternative heuristics, when tested with scatter plots of 5 to 20 data points. This previous study, however, does not provide a best exponent as an outcome of an inverse optimization analysis, and instead, compared a few different methods that are not on a continuum.

There are studies reporting that there is a central tendency (Hollingworth, 1910) when humans perceive a dataset. Many of our subjects indeed responded to our optional questionnaire that their strategy was to “put the line at the center of the distribution” when they performed the regression or classification task, but the best-describing loss functions varied to a large degree even between these subjects. The definition of “central” differs from study to study when referring to central tendency. For example, the mean, median, and mode of the data could all reasonably be defined to be the center of a distribution. The best-describing parameters we found from the experiment could be interpreted in terms of these statistical quantities of the distribution, because finding the median and mean are mathematically equivalent to finding a regression line that minimizes the sum of absolute regression error and error squared. Using this equivalent description, our inverse optimization result on the regression task is that, people on average chose a regression line near mean for high sparsity data, near median for medium sparsity data, and between median and mode for low sparsity datasets. While “central” is even less well-defined for two distributions for binary classification tasks, our result from classification shows that there was a similar shift with the sparsity of the data: the average decision boundary moved towards the skew-symmetric distribution as sparsity decreased. Our results provide some insights into central tendency theory, suggesting that “central” could be highly context-dependent and highly variable across people.

Traditional computational algorithms are normally not designed to change their strategies based on the size of the data, but we observed noticeable shifts in average strategy when humans perform regression and classification. Potential explanations for the shift we observed include: when more data points were presented, (1) people could infer the underlying structure of data better, and thus made different judgments on the importance of each data point; (2) people may perceive the nature of the dataset differently, e.g., less sparse data (dense data points) might be perceived as a contiguous area, whereas sparse data would be perceived as a set of individual dots; (3) it becomes more challenging to consider individual data simply because there are a lot of them, thus people switched to different heuristics. The third point seemed to be more apparent for the regression task, where subjects took longer times to complete a high-sparsity task that had fewer dots than a lower-sparsity task that had more dots. These shifts in behaviors, both in terms of the decision itself and the time taken to make the decision, provide insights into how humans perceive patterns differently in data of different sizes.

Commonly used loss functions for regression or classification algorithms are mostly convex, so that the loss function is smooth and there is a single global minimum. For example, the sum of distances raised to the exponent smaller or equal to 1 could have a non-smooth loss function, and could result in multiple local and global minima (figure 4), and thus are not desirable for machine learning algorithms. However, many subjects chose regression lines and decision boundaries in the range that was only reached by loss function models that were non-convex, at least among the loss function models we considered in this study. Another limitation that comes from using only convex loss functions is that the contributions from the outliers are bound to be bigger than a certain value. Loss functions that are relatively robust to outliers, such as Huber loss and log-cosh loss, have smaller increases in slope at the extremes than near zero, but only to the degree that the overall loss function is still convex. Although convexity and other traits are desirable for machine learning algorithms, it is conceivable that computational and mathematical simplicities are not as strictly required for humans. Therefore, developing a machine learning algorithm that closely mimics human perception of patterns may necessitate the use of loss functions that are not conventionally used.

Unlike computational optimization, “noise” can be a factor in how humans actually perform pattern recognition, even when they have a consistent criterion (which is not necessarily guaranteed). Our experimental results exhibited high intra-subject variability, and noise could be one possible explanation. Noise in the human performance (Kahneman et al., 2021), arising from various sources such as limitations on calculation precision, inaccurate motor execution while indicating the selection, and even effects of optical illusion, were hoped to be canceled out if we increase the experimental sample size and look for an average behavior across samples. However, loss functions of some types could result in the ill-posedness of the inverse optimization problem, which may not be rectified by simply adding more samples to the analysis. Consider the case when a loss function is very flat near the minima, or when it has multiple local minima that have loss values close enough (or even exactly the same) to the global minimum (e.g., figure 4). Even if a subject is consistently using the specific loss function, they may still make a final decision that is not exactly at the global minimum, but at other reasonable alternatives, like, at one of the local minima. In this case, the loss function could falsely appear to be an inadequate descriptor of subjects’ decisions. We partially addressed this issue by considering multiple global minima locations for p<=1 and by assuming that the location that was closer to what subjects selected was the location they were aiming for. However, this assumption may have given p<=1 an unfair advantage against p>1. In addition, this still does not address the case of humans selecting a location that is close enough to the global minimum but numerically not identical. Further studies would be needed to improve inverse optimization analysis when the proximity between the observed decision and the specific optimization criterion is an ill-behaved function like this.

There was high inter-subject variability in our experimental results. Sometimes patterns in the visually presented data seem obvious (Wertheimer, 1938) and can be easily agreed between different people. However, the high inter- and intra-subject variability that we observed on relatively simple tasks leads to a suspicion that there might be no single commonsensical “ground truth” to many of the regression and classification problems. As some authors point out, (Von Luxburg et al., 2012), some pattern recognition may need to be viewed more like art than science.

There were some limitations to our current study. We imposed the patterns to be one-dimensional, but higher dimensional problems may result in different results. We also cannot test overfitting with one-dimensional problems, but in future studies, we could consider introducing a higher degree of freedom to see what people do. Most of our subjects had a science and engineering background, and had some previous experience with regression techniques, while fewer had an experience with classification techniques. Our optional questionnaire shows that about half of the subjects could perform a regression method they learned, and half of the subjects at least learned about it in the past, whereas about half of the subjects never heard about classification algorithm and very few subjects responded that they could perform a classification method. As shown in a previous study (Gillan, 2020), people with statistics training seem to use a different method to perform regression than people without such training, and we also expect that our result is dependent on the subject group. Results might also be dependent on the instruction, and specific details of the test such as the scaling of the task interface on the computer screen (as it was reported in a previous study that the scaling of the graph affected subjects’ decisions (Cleveland et al., 1982)), color and size of the dots, and the order of the trials. We applied the inverse optimization analysis to loss functions that simply add the contributions from each data point, but it would be also interesting to test other forms of loss functions.

Statistics of the sample data are different from the statistics of the PDF from which the dots were drawn from, thus regression lines or decision boundaries that are calculated based on sample data are also different from the calculations based on the PDF (figure 2). These discrepancies are bigger when we draw a smaller number of dots. Since the underlying distribution could become more obvious when people have seen the dots of lower sparsity, and this could be used as a priori knowledge when they perform the task on high sparsity data, we tested most of the subjects in partially randomized orders, such that they performed higher sparsity tasks first. However, when we compare the subsets of subjects (*N*_subjects_ = 7) who were tested in a fully randomized order to the rest of the subjects, we found no evidence that the different randomization had a significant effect (two-sample t-test had *p* valu of 0.15 for the regression task, and 0.95, 0.80, 0.46 for classification task using rectified error, log power, log margin loss function). However, given the high variability of the results, order effects could be studied with a bigger sample size.

Another way subjects may apply a priori knowledge while performing the tasks could be by estimating where the center of the dataset would be, based on the experience of past trials. The expected location of the center of the datasets was roughly in the middle of the visual area, as we randomized the vertical location of the scatter plot on the testing interface. Thus, if humans are updating a Bayesian prior during the experiment to make decisions based on a mixture of their estimated prior and the current dataset, we might expect systematic deviations toward the center of the visual area. However, we do not see this deviation in the data (figure 10).

**Figure 10:**
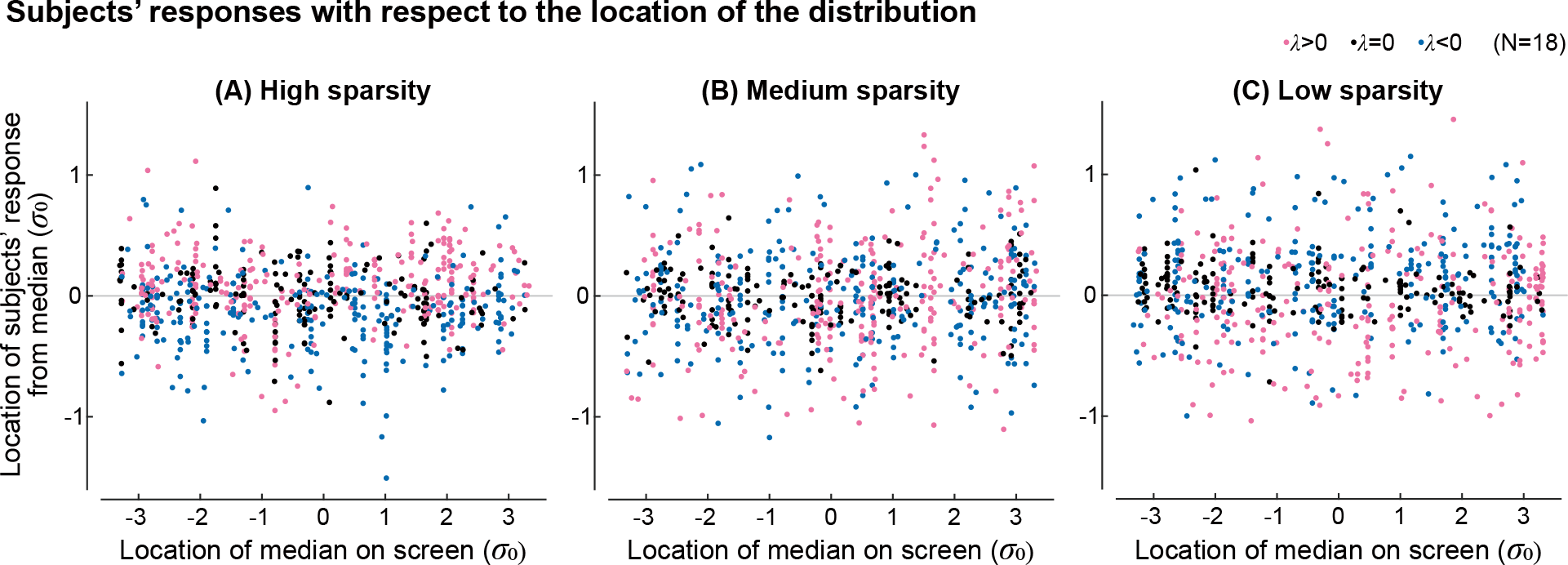
Error dependence on location on screen. The error with respect to the median or any other *L*_*p*_ optimum does not depend on the location of the distribution’s median on the screen.

We designed datasets to study what loss functions humans optimize for when they perform regression and classification, and obtained loss functions that were on average best descriptors of subjects’ responses. However, the result was not as simple as simply reporting what was “the” loss function humans optimize. Instead, we learned that the loss function could be context-dependent, and that there are high subject variabilities when humans perform visual pattern recognition tasks. It would be interesting to see what are the other factors that contribute to the human inconsistency, as well as measuring the inconsistency itself more closely, for example by repeating the same test again. More in-depth investigation of how humans perform an individual trial could also provide useful insights. Eye tracking could show what points each subject considered while performing the task and may especially provide insights into outlier processing. We could also ask people to speak out what their thoughts are while perceiving the dots. Another direction for a future study could be to use specially designed probability distributions where dots are drawn from, so that specific hypotheses could be tested. The way data is presented could be also altered. For example, one could present a target line and ask people to move the set of dots to best hit the target line. One could present some dots each moment and refresh the dot locations with some time interval, investigating the history effect on pattern perception. These tests could be also integrated with motor control tasks, for example, humans performing aiming, reaching, or catching tasks, while target objects to interact with have some probability distribution in terms of their position or velocity.

We obtained average loss functions that human subjects use when they perform regression and classification tasks, while observing a large variability. This observation would contribute to the understanding of human visual perception and motor control using visual perception, and would provide insights into developing machine learning techniques. Understanding how humans perform such pattern recognition tasks can be useful as ground truth to train machine learning algorithms aimed at reproducing human intelligence (Foncubierta Rodríguez & Müller, 2012), as an inspiration to develop new algorithms, since humans seem capable of performing complicated pattern recognition tasks, to identify common mistakes humans make and systematic biases that humans have, so as to educate people (Tschandl et al., 2019; Kahneman, 2011), and more broadly, to study how human nervous system functions (Körding & Wolpert, 2004a; Schwab & Nusbaum, 2013; Caelli et al., 1987).

## Author Contributions

HXR and MS conceived the study. HXR designed and performed the experiments, and analyzed the data. HXR wrote the first draft of the manuscript and both authors revised the manuscript.

## Acknowledgments

HXR was supported by the U. Calgary Department of Biomedical Engineering. MS was supported in part by NIH-R01GM135923-01 and NSF SCH grant 2014506.

## References

Adelchi Azzalini. A class of distributions which includes the normal ones. Scandinavian journal of statistics, pp. 171–178, 1985.

Sheldon Baron and David L Kleinman. The human as an optimal controller and information processor. IEEE Transactions on Man-Machine Systems, 10(1):9–17, 1969.

Christopher M Bishop and Nasser M Nasrabadi. Pattern recognition and machine learning, volume 4. Springer, 2006.

Terry Caelli, Ingo Rentschler, and W Scheidler. Visual pattern recognition in humans: I. evidence for adaptive filtering. Biological Cybernetics, 57(4-5):233–240, 1987.

William S Cleveland, Persi Diaconis, and Robert McGill. Variables on scatterplots look more highly correlated when the scales are increased. Science, 216(4550):1138–1141, 1982.

Michael Correll and Jeffrey Heer. Regression by eye: Estimating trends in bivariate visualizations. In Proceedings of the 2017 CHI Conference on Human Factors in Computing Systems, pp. 1387–1396, 2017.

Keith Davids, A Mark Williams, and John G Williams. Visual perception and action in sport. Routledge, 2005.

Winand H Dittrich. Action categories and the perception of biological motion. Perception, 22(1):15–22, 1993.

Harris Drucker, Christopher J Burges, Linda Kaufman, Alex Smola, and Vladimir Vapnik. Support vector regression machines. Advances in neural information processing systems, 9, 1996.

Antonio Foncubierta Rodríguez and Henning Müller. Ground truth generation in medical imaging: a crowdsourcing-based iterative approach. In Proceedings of the ACM multimedia 2012 workshop on Crowd-sourcing for multimedia, pp. 9–14, 2012.

Douglas J Gillan. Fitting regression lines to scatterplots: The role of perceptual heuristics. In Proceedings of the Human Factors and Ergonomics Society Annual Meeting, volume 64, pp. 1650–1654. SAGE Publications Sage CA: Los Angeles, CA, 2020.

Leon Glass and Rafael Pérez. Perception of random dot interference patterns. Nature, 246(5432):360–362, 1973.

Leon Glass and Eugene Switkes. Pattern recognition in humans: Correlations which cannot be perceived. Perception, 5(1):67–72, 1976.

James Gopsill, Mark Goudswaard, David Jones, and Ben Hicks. Capturing mathematical and human perceptions of shape and form through machine learning. Proceedings of the Design Society, 1:591–600, 2021.

Arnulf BA Graf, Felix A Wichmann, Heinrich H Bülthoff, and Bernhard Schölkopf. Classification of faces in man and machine. Neural Computation, 18(1):143–165, 2006.

Norbert Henze. A probabilistic representation of the’skew-normal’distribution. Scandinavian journal of statistics, pp. 271–275, 1986.

Brian L Hills. Vision, visibility, and perception in driving. Perception, 9(2):183–216, 1980.

Harry Levi Hollingworth. The central tendency of judgment. The Journal of Philosophy, Psychology and Scientific Methods, 7(17):461–469, 1910.

Daniel Kahneman. Thinking, fast and slow. Macmillan, 2011.

Daniel Kahneman, Olivier Sibony, and Cass R Sunstein. Noise: A flaw in human judgment. Little, Brown, 2021.

Daniel A Keim, Florian Mansmann, Jörn Schneidewind, and Hartmut Ziegler. Challenges in visual data analysis. In Tenth International Conference on Information Visualisation (IV’06), pp. 9–16. IEEE, 2006.

Konrad P Körding and Daniel M Wolpert. Bayesian integration in sensorimotor learning. Nature, 427(6971): 244–247, 2004a.

Konrad Paul Körding and Daniel M Wolpert. The loss function of sensorimotor learning. Proceedings of the National Academy of Sciences, 101(26):9839–9842, 2004b.

Stephan Lewandowsky and Ian Spence. The perception of statistical graphs. Sociological Methods & Research, 18(2-3):200–242, 1989.

C Karen Liu, Aaron Hertzmann, and Zoran Popović. Learning physics-based motion style with nonlinear inverse optimization. ACM Transactions on Graphics (TOG), 24(3):1071–1081, 2005.

David G Luenberger. Optimization by vector space methods. John Wiley & Sons, 1997.

Hamed Masnadi-Shirazi and Nuno Vasconcelos. A view of margin losses as regularizers of probability estimates. The Journal of Machine Learning Research, 16(1):2751–2795, 2015.

Jeanne Moore. Data visualization in support of executive decision making. Interdisciplinary Journal of Information, Knowledge, and Management, 12:125, 2017.

Frederick Mosteller, Andrew F Siegel, Edward Trapido, and Cleo Youtz. Eye fitting straight lines. The American Statistician, 35(3):150–152, 1981.

Stephen E Palmer. Visual perception and world knowledge: Notes on a model of sensory-cognitive interaction. Explorations in cognition, pp. 279–307, 1975.

Eileen C Schwab and Howard C Nusbaum. Pattern recognition by humans and machines: speech perception, volume 1. Academic Press, 2013.

Michael Sedlmair, Andrada Tatu, Tamara Munzner, and Melanie Tory. A taxonomy of visual cluster separation factors. In Computer Graphics Forum, volume 31, pp. 1335–1344. Wiley Online Library, 2012.

John E Shore and Robert M Gray. Minimum cross-entropy pattern classification and cluster analysis. IEEE Transactions on Pattern Analysis and Machine Intelligence, (1):11–17, 1982.

Manoj Srinivasan. Optimal speeds for walking and running, and walking on a moving walkway. Chaos: An Interdisciplinary Journal of Nonlinear Science, 19(2), 2009.

Albert Tarantola. Inverse problem theory and methods for model parameter estimation. SIAM, 2005.

Emanuel Todorov and Michael I Jordan. Optimal feedback control as a theory of motor coordination. Nature neuroscience, 5(11):1226–1235, 2002.

Philipp Tschandl, Noel Codella, Bengü Nisa Akay, Giuseppe Argenziano, Ralph P Braun, Horacio Cabo, David Gutman, Allan Halpern, Brian Helba, Rainer Hofmann-Wellenhof, et al. Comparison of the accuracy of human readers versus machine-learning algorithms for pigmented skin lesion classification: an open, web-based, international, diagnostic study. The lancet oncology, 20(7):938–947, 2019.

Ulrike Von Luxburg, Robert C Williamson, and Isabelle Guyon. Clustering: Science or art? In Proceedings of ICML workshop on unsupervised and transfer learning, pp. 65–79. JMLR Workshop and Conference Proceedings, 2012.

Howard Wainer and David Thissen. On the robustness of a class of naive estimators. Applied Psychological Measurement, 3(4):543–551, 1979.

Qi Wang, Yue Ma, Kun Zhao, and Yingjie Tian. A comprehensive survey of loss functions in machine learning. Annals of Data Science, 9(2):187–212, 2022.

Max Wertheimer. Laws of organization in perceptual forms. 1938.

Bang Wong. Color blindness. nature methods, 8(6):441, 2011.

